# Identification of bacterial candidates that promote the growth of the seagrass *Zostera marina*

**DOI:** 10.64898/2026.03.19.712741

**Authors:** Diane-Marie Brache-Smith, Jacquelyn Badillo, Saray Maeda, Maggie Sogin

## Abstract

**Background:** Globally, seagrass ecosystems are threatened by anthropogenic activities that are leading to increased levels of eutrophication, coastal pollution and thermal conditions. Consequently, there is a growing need to develop new approaches that work to mitigate these stressors and enhance restoration efforts in seagrass meadows. One promising strategy is to identify, isolate and characterize microbial consortia that are likely to support seagrass productivity. However, our current understanding of key microbial functions that support plant growth in marine systems is limited. Based on evidence from terrestrial plant-microbe systems, seagrass-associated bacteria are expected to provide the plant with nitrogen and phosphorus resources while detoxifying sulfur and producing phytohormones. Here, we sequenced 61 bacterial cultures isolated from the rhizosphere, rhizoplane, and endosphere of the seagrass, *Zostera marina* to identify a consortium of six putative plant growth promoting (PGP) candidates.

**Results:** Our cultivation approach using plant-based media allowed us to isolate 201 bacteria from *Z. marina*, which reflected 18% of the total microbial diversity of the starting inoculum. Genomic and phenotypic analyses of the 61-sequenced pure-cultures revealed that most of the sequenced taxa were able to mobilize nitrogen primarily through catabolic pathways, including denitrification (51%), dissimilatory nitrate reduction to ammonia (71%), and C-N bond cleavage (83%). Six of the isolates, which represent new lineages of *Agarivorans*, coded for the nitrogenase gene cassette. Additionally, 52% of the genomes had genes for sulfur and/or thiosulfate oxidation, 88.5% for phosphorus solubilization, and 60.5% for IAA production. Genomic analysis also revealed that some pathways, including denitrification and dissimilatory nitrite to ammonia DNRA, required cross-species cooperation as no one taxa contained all the genes needed to complete these metabolic pathways. Based on draft genome models and results from phenotypic assays, isolates *Streptomyces sp*. (Iso23 and Iso384), *Mesobacillus sp* (Iso127), *Roseibuim sp.* (Iso195), *Peribacillus sp.* (Iso49), and *Agarivorans sp.* (Iso311) represent a minimal microbial community that is likely to promote seagrass growth and enhance restoration efforts.

**Conclusion:** Our work provides a detailed genomic and phenotypic analysis of bacteria isolated from *Z. marina* and identifies a minimal microbial community with complementary PGP traits. Isolating, identifying and characterizing bacteria that promote seagrass growth is critical towards enhancing restoration efforts of seagrass meadows.

## BACKGROUND

Seagrasses are among the most productive ecosystems on Earth. While only occupying less then 0.2% of the world’s oceans, they provide key habitat for marine fisheries, protect coastlines from wave energy and erosion and store up to 20% of marine organic carbon^1,2^. Unfortunately, these ecosystems, valued at over cost of $213 billion (2020 US dollars) to the global economy, are experiencing rapid decline^3^. Global oceans are losing seagrass meadows at a rate of 7% per year, which is resulting in a net release of 1,154 Tg of carbon (C) back to the atmosphere^3,4^. Declines in water quality, coastal eutrophication and shifts in sediment biogeochemistry are limiting the success of seed-based and plant transplantation strategies aimed at restoring meadows^5–7^. To conserve seagrasses worldwide, there is a need to develop new techniques to supported seagrass growth under these challenging environmental conditions.

The plant growth promoting (PGP) bacteria and the related emerging field of wildlife probiotics are predicted to enhance plant and animal health across biological systems^8,9^. For example, an engineered synthetic community of rhizosphere microorganisms effectively promoted the growth of sorghum in both laboratory and field settings^10^. In tomatoes, inoculation with bacterial strains isolated from the plant’s microbiome with PGP traits increased plant growth and root development^11^. Similar microbiome manipulations through the addition of beneficial bacteria shows promise for enhancing host resilience in marine systems in general^12^, and seagrasses in particular. For example, in reef-building corals a beneficial strain of *Pseudoalteromonas* worked to reduce disease progression in colonies experiencing stony coral tissue loss disease^13^. Beneficial microbes also protect corals from mortality during a thermal stress^14^. In seagrasses, the inoculation of two bacteria taxa increased photosynthetic performance of the seagrass *Thalassia hemprichii*^15^. Despite this early success in developing candidates for plant growth promotion for marine plants, we lack culture representatives with known functions in supporting seagrass growth.

Emerging evidence indicates that seagrasses host diverse microbial consortia in their roots and rhizosphere sediments that play multiple roles in host health, including supporting plant growth^16–18^. For example, seagrasses associate with sulfur-oxidizing bacteria that reduce the toxic effects of sulfide build up^19,20^, and with nitrogen-fixing bacteria that provide the plant with nitrogen resources^21^. It is less clear if and how members of the seagrass microbiome support plant acquisition of phosphorus and produce hormones, including indole-3-acetic acid (IAA). Yet, in terrestrial systems, these functions are key drivers of plant growth^22^. A key barrier in determining the capacity of seagrass associated bacteria in supporting plant growth is that many members of the seagrass microbiome are recalcitrant to cultivation and isolation approaches. This is likely a consequence of using standard media conditions to isolate host-associated microorganisms^23^. To overcome this limitation, one approach is to develop enrichment media containing host-derived factors, which may select for ecologically relevant, plant-associated taxa, with beneficial phenotypes^23,24^.

Here, we isolated, characterized and identified putative PGP bacteria for the seagrass, *Zostera marina*. We analyzed the genomes of 61 pure-culture bacterial strains with a focus on taxa that tested positive for plant growth promotion traits, including N-mobilization, S-reduction, S-oxidation, P-solubilization and the production of IAA. We then applied genome-scale metabolic models to identify a minimal community of microbial taxa that had complementary functions. Using our approach, we proposed a community of microbial taxa that are predicted to promote seagrass growth.

## METHODS

### Cultivation, identification and characterization of isolates

To cultivate diverse consortium of bacteria from the rhizosphere, rhizoplane and endosphere of the seagrass, *Z. marina*, we developed a cultivation media where complex metabolite extracts from seagrass tissues represented the sole carbon source (**Supplementary Figure 1**). Our approach takes advantage of host derived factors, for example plant produced metabolites and nutrients, to select for microbial species that require these compounds for growth^23,25^ . To generate the plant-based cultivation media (PB), we collected *Z. marina* roots and rhizomes from Bodega Bay, California, USA (38.333446, −123.058785). The plant tissues were rinsed with sterile seawater and subsequently ground into a fine powder under liquid nitrogen. To make the PB media, 5 g of the ground plant material was added to 45 mL of sterile minimal artificial seawater (SASW), along with 32 mm ceramic beads, and incubated overnight at 22°C at 200 rpm. Following incubation, the slurry was filter-sterilized (0.22 µm) to remove bacteria and plant debris. To prepare the PB plates, 50 mL sterilized slurry was added to 950 mL SASW with 15 g noble agar.

To isolate bacteria from *Z. marina* rhizosphere, rhizoplane and endosphere, we collected replicate (n=3) individuals of *Z. marina* during the plant’s growing season from Bodega Bay, California, USA (38.333446, −123.058785). First, excess non-rhizosphere sediments were gently removed from the plants, and the rhizomes were trimmed to 3 cm. To generate the rhizosphere inoculum, the roots and rhizomes were submerged in 10 mL of SASW and vortexed to collect the rhizosphere sediments. After which, the plant roots and rhizomes were washed in SASW and transferred to a fresh 50 mL Falcon tube containing 10 mL of SASW. To obtain the rhizoplane inoculum, the roots were sonicated at room temperature (3 x 30 s). In between sonication rounds, the samples were placed on ice to avoid overheating the sample. The roots and rhizomes were then surface sterilized in 99% ethanol and rinsed three times in 10 mL of SASW to remove remaining epiphytes. To confirm the success of the surface sterilization, rhizomes were briefly vortexed in 10 mL of SASW and the resulting slurry was plated on Marine Broth (MB) 2216 (Sigma Aldrich) and PB agar plates to check for growth. Finally, to obtain the endosphere inoculum, the sterilized roots and rhizomes were macerated and transferred to a Falcon tube containing 10 mL of SASW, and sonicated (3 x 30 s) to release endophytic bacteria into the cell slurry. All inoculum samples were stored on ice prior to plating for cultivation. 100 µL of each serial dilution up to 10^2^ from each inoculum was plated on PB plates and incubated either at 14°C or 22°C for 30 days. Individual colonies (n=10-16 per plate) were picked and purified on either to PB agar or MB plates. Purified isolates were stored at −80°C on 45% glycerol until revived for DNA extraction.

To identify *Z. marina* isolates, a single colony (n=201) was picked and crush in 100 µl 50% dimethyl sulfoxide at 60°C for 20 minutes. We preformed colony PCR of the 16S rRNA gene using the universal primers GM3F (5’-AGAGTTTGATCMTGGC-3’) and GM4R (5’-TACCTTGTTACGACTT-3’)^26^ under the following cycling conditions: 3 min at 98 °C; 30 cycles of 30 s at 98 °C, 30 s at 55 °C, 1 min at 72 °C; 2 min at 72 °C. PCR reactions were conducted using Q5 High-Fidelity taq-DNAp Polymerase (NEB) and 2% dimethyl sulfoxide. Sequencing reactions and PCR clean-up were conducted by Molecular Cloning Laboratories (MCLAB) (South San Francisco, CA), using the ABI 3730XL sequencer. The resulting sequences were quality trimmed using the *TrimEnds* function in Geneious Prime v2024.0.4^27^ with an error probability limit of 0.05 to an average of 542 bp. To identify each isolate, we used BLASTn^28^ to compare the cleaned sequence to the GTDB r226 16S rRNA gene reference database^29^.

We screened taxonomically distinct isolates in vitro to quantify their activity for four plant-growth-promoting traits. First, to determine if each isolate mobilized nitrogen, we grew each isolate on Jensen’s nitrogen-free medium with Bromothymol blue as a color indicator^30^. Phosphorus solubilization was screened by plating each isolate on Pikovskaya’s Agar^31^. Sulfur oxidation or reduction was determined by growing each isolate on thiosulfate media with 0.5% phenol red indicator^32^. Finally, We measured IAA production using the Salkowski Assay, where overnight cultures of each isolate were grown in tryptophan-supplemented on MB^33^. The supernatant was treated with Salkowski reagent for colorimetric quantification. Isolates (n=61) that tested for two or more putative PGP traits were targeted for whole genome sequencing (WGS).

### 16S rRNA amplicon sequencing of the inoculum

To characterize the microbiome composition of each inoculum sample using culture independent techniques, we mixed 500 µL of the undiluted homogenate with an equal volume of Zymo DNA/RNA Shield at the time of plating. Genomic DNA was extracted from 300 μL of reserved homogenate following the Cetrimonium bromide (CTAB) extraction protocol^34^ with the following modifications: samples were incubated with CTAB lysis buffer (2.5 M NaCl, 2% CTAB, 0.1% PVPP) at 65°C for 20 min, bead-beaten at 5.5 m/s for 45 s, then purified by phenol:chloroform extraction and ethanol precipitation. As a negative control, 3 L of sterile artificial seawater (SASW) used for dilutions was filtered and extracted following the same protocol.

To assess how well the bacterial isolates represent the community of *Z. marina* rhizosphere, rhizoplane and endosphere, we preformed amplicon sequencing of the full-length region of the 16S rRNA gene. Genomic DNA samples were amplified using Q5 High-Fidelity 2× Master Mix (New England Biolabs) with peptide nucleic acid clamps universal pPNA sequence (5’-GGCTCAACCCTGGACAG-3′) and universal mPNA clamps (5’-GGCA AGTGTTCTTCGGA-3′)^35^ added to a final concentration of 0.5 μM to block amplification of host plastid and mitochondrial sequences. The 16S rRNA gene were amplified using GM3F (5’-AGAGTTTGATCMTGGC-3’) and GM4R (5’-TACCTTGTTACGACTT-3’)^36^. PCR amplification was performed under the following cycling conditions: initial denaturation at 98°C for 30 s; 35 cycles of 98°C for 10 s, 56°C for 30 s, and 72°C for 1 min; final extension at 72°C for 2 min. PCR products were purified using solid-phase reversible immobilization (SPRI) magnetic beads and subsequently prepared for sequencing using the Oxford Nanopore Technology (ONT) sequencing using the SQK-NBD114 Native Barcoding Kit with Short Fragment Buffer to prevent loss of amplicons. Sequencing was performed on the PromethION 2 Solo platform using a FLO-PRO114M flow cell (R10.4.1 chemistry) at SeqCoast Genomics (New Hampshire, USA). Base calling was performed using Dorado v7.4.12^37^ in super-accurate basecalling mode and barcode trimming enabled in MinKNOW (v24.06.10). To calculate ASVs, we used NanoASV v1.2.2^38^ with a minimum quality score threshold of Q20 to calculate amplicon sequence variants (ASVs) from the ONT data. The R-based quality filtering step was disabled (--no-r-cleaning) to retain ONT-specific quality metrics. Taxonomic classification was performed by comparing ASV sequences to the GTDB r226 16S rRNA database^29^. The resulting phyloseq object was decontaminated using prevalence-based filtering using control samples (decontam), and taxonomy was cleaned by removing unclassifiable clusters, correcting known misannotations microViz0.12.7^39^, removing GTDB sub-clade suffixes, and filtering singleton ASVs (total abundance < 2). The resulting sequences of all 2,378 ASVs sequences were aligned using MAFFT v7.525^40^ (--adjustdirectionaccurately), and a maximum-likelihood phylogenetic tree was constructed using IQ-TREE v2.1.36^41^ using GTR+G tree Model and 5000 ultrafast bootstrap replicates. After generating the base tree, 16S rRNA gene sequences were retrieved using Barrnap v0.9^42^ from the 61 whole-genome assemblies (see **DNA extraction, sequencing and assembly of isolate genomes**). These extracted sequences, along with 16S rRNA gene sequences from Sanger-sequenced colony PCR products (n=140), were phylogenetically placed onto the reference ASV-based tree using App-SpaM^43^. The resulting tree was visualized using iTOL v7.2.28^44^.

### DNA extraction, sequencing and assembly of isolate genomes

Genomic DNA was extracted from freshly grown isolates (OD_600_ >1) using the Quick-DNA Miniprep Plus Kit (ZymoResearch, Irvine, CA), with a lysozyme pre-treatment for Gram-positive species. Genome sequencing was performed by Plasmidsaurus (Eugene, OR) using Nanopore sequencing with the ONT Ligation sequencing kit v.14 (SQK-LSK114) without size selection or fragmentation according to the manufacturer’s instructions. Libraries were sequenced on a PromethION 24 device using R10.4.1 flow cell chemistry. Base calling was completed using Guppy v.6.1.5. Raw reads were filtered using Filtlong v.0.2.1^45^ using the flags –min length 1000 and –keep percent 90 to remove reads shorter than 1,000 bp and retain the top 90% of reads based on quality. Consensus long read assemblies were generated with Autocycler v.0.2.1^46^ using its default set of assemblers (CANU, Flye, miniasm, necat, NextDenovo, and Raven). The circular assembly graph was manually inspected using Bandage^47^ and subsequently reoriented to the origin of replication using Dnaapler v.1.2.0^48^. The completeness and contamination of each genome was assessed with CheckM v1.2.3^49^.

### Genome annotation and identification of genomic features, pathways and genes of interest

Genome assemblies were annotated using Prokka v1.14.6^50^ with the (--kingdom Bacteria) flag and default parameters, including gene prediction (--addgenes), mRNA annotation (--addmrna), and non-coding RNA identification via Rfam (--rfam). Functional annotation and ortholog assignment were performed using eggNOG-mapper v2.1.12^51^, which aligned Prokka-predicted protein sequences against the eggNOG v5.0.2 database using DIAMOND v2.1.11^52^. KEGG orthologs (KO) and functional predictions were extracted from the eggNOG-mapper output for downstream pathway analysis. Taxonomic classifications were assigned using the Genome Taxonomy Database Toolkit (GTDB-Tk v 2.4.0^53^) with the classify_wf workflow against GTDB release r220^54^. To assess genomic relatedness among isolates, pairwise Average Nucleotide Identity (ANI) values were calculated using FastANI v1.34^55^. A phylogenomic tree was constructed from all isolate genomes using GToTree v1.8.8^56^ with the bacterial single-copy gene set (74 target genes).

### Minimal community selection using genome-scale metabolic modeling

Genome-scale metabolic network (GSMN) reconstruction and minimal community analysis were performed using the Metage2Metabo v1.6.1 (M2M Mincom) pipelinee^57^. GSMNs were reconstructed for 61 bacterial isolates using the recon module, which automates Pathway Tools v29.0^58^ with the MetaCyc Metabolic Pathway Database^59^ as a reference. Annotated genomes in GenBank format served as input, and the resulting GSMNs were exported in SBML files. Seed metabolites (n=73) simulated the *Z. marina* rhizosphere^60,61^, including nitrogen fixation substrates, phosphorus sources, sulfur compounds, and essential cofactors (**Supplementary Table 1**)^62^. Target metabolites (n=30) represented plant growth-promoting compounds: nitrogen assimilation products, phosphorus mobilization products, sulfur detoxification endpoints, auxin (indole-3-acetic acid) and biosynthetic intermediates, polyamines, and siderophores (**Supplementary Table 1**). The M2M Mincom pipeline identified minimal bacterial communities and classified bacteria as essential “symbionts” (present in all solutions) or alternative “symbionts” (present in at least one solution). The M2M cscope pipeline was used to identify community producible metabolites of the minimal community consortia.

## RESULTS

### Diversity of bacteria cultivated from *Z. marina*

We recovered an average of 15,676 ± 13,393 reads per sample from the 16S rRNA amplicon libraries from inoculate samples collected from *Z. marina’s* rhizosphere, rhizoplane and endosphere. Overall, we calculated a total of 2378 ASVs across the entire dataset with 1111 ASVs from the rhizosphere, 1446 ASVs from rhizoplane and 1540 ASVs from the endosphere. The vast majority (79.5%) of the ASVs were classified to the genus level. Gammaproteobacteria (20.17 %), Alphaproteobacteria (17.39 %), Bacteroidia (11.84 %), and Desulfobulbia (7.35 %) bacterial classes dominated the relative abundance of the bacterial community across all compartments, while Actinomycetes (2.35%) and Bacilli (2.06%) had lower relative abundances.

In total, we isolated 201 bacteria cultures from *Z. marina*. Phylogenetic placement of the isolates’ 16S rRNA sequences onto the 16S rRNA gene tree showed that cultured taxa were distributed across the phylogeny (**Fig. 1**). Specifically, we were able to isolate representatives from Gammaproteobacteria (n=115), Bacilli (n=55), Actinomycetes (n=27), Alphaproteobacteria (n=3), and Bacteroidia (n=1) groups (**Fig. 1, Supplementary Fig. 2b**). Due to our cultivation approach, we were unable to isolate members of Acidimicrobiia, Anaerolineae, Campylobacteria, Clostridia, Desulfobacteria, Desulfobulbia, and Planctomycetia, which are largely predicted to be anaerobic. All four of the classes that we were able to isolate members from were recovered in *Z. marina’s* rhizosphere, rhizoplane and endosphere. The isolates spanned 24 families, 42 genera and 86 species (**Supplementary Table 2**), and were largely dominated by Bacillaceae (n= 46), Celerinatantimonadaceae (n=30), Marinomonadaceae (n=28), Vibrionaceae (n=20), Alteromonadaceae (n=13), and Cellvibrionaceae (n=13). There were 18 additional families that had fewer than 10 isolates per group. In total, we estimate that we were able to culture members from approximately 18% of the families represented in the 16S rRNA amplicon dataset (**Supplementary Fig. 2a, b**).

**Fig 1.**
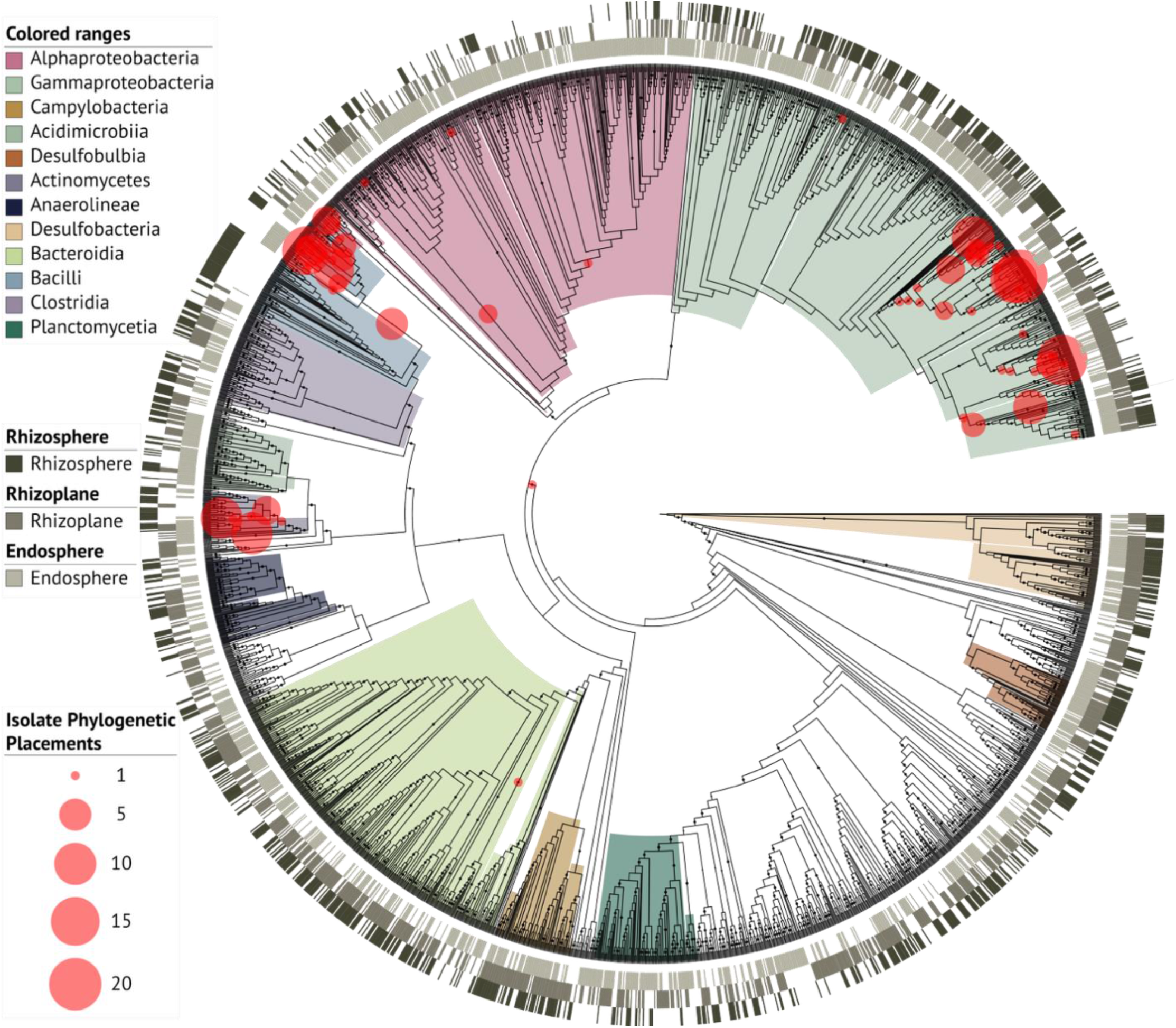
Isolate placement (red circles) on the phylogenetic tree of the ASV sequences generated from *Z. marina* endosphere, rhizoplane and rhizosphere inoculum samples show that the culture collection consisted of lineages primarily from the Gammaproteobacteria, Alphaproteobacteria, Bacilli and Actinomycetes groups. Red circles indicate phylogenetic placements of the 16S rRNA sequence from cultured isolates determined using App-SpaM^43^. Colored strips adjacent to ASV branches indicate the plant compartment where each ASV was detected while phyla are indicted by color ranges on the tree. Bootstrap support >98% are shown. Size of the red circles represents the number of isolates placed on the tree at each node.

Using the taxonomic diversity of the 16S rRNA Sanger sequences and assay results as a guide (**Supplementary Fig. 3**), we selected 61 out of the 201 isolates for WGS. These isolates included representatives of 28 genera, including *Vibrio* (n=7), *Agarivorans* (n=6)*, Micrococcus* (n=6), *Nocardiopsis* (n=6*), Anaerobacillus* (n=5), *Streptomyces* (n=5), and *Marinomonas* (n=5) (**Fig. 2; Supplementary Fig. 4**). Based on GTDB-tk classification of the WGS, 17 of the 61 isolates could only be classified to the genus level, suggesting that they are representative of new bacterial species. These likely novel species included members of *Agarivorans*, *Isoptericola*, *Marinomonas*, *Nocardiopsis*, *Roseibium*, *Rossellomorea*, *Shewanella*, and *Solibacillus* genera.

**Fig 2.**
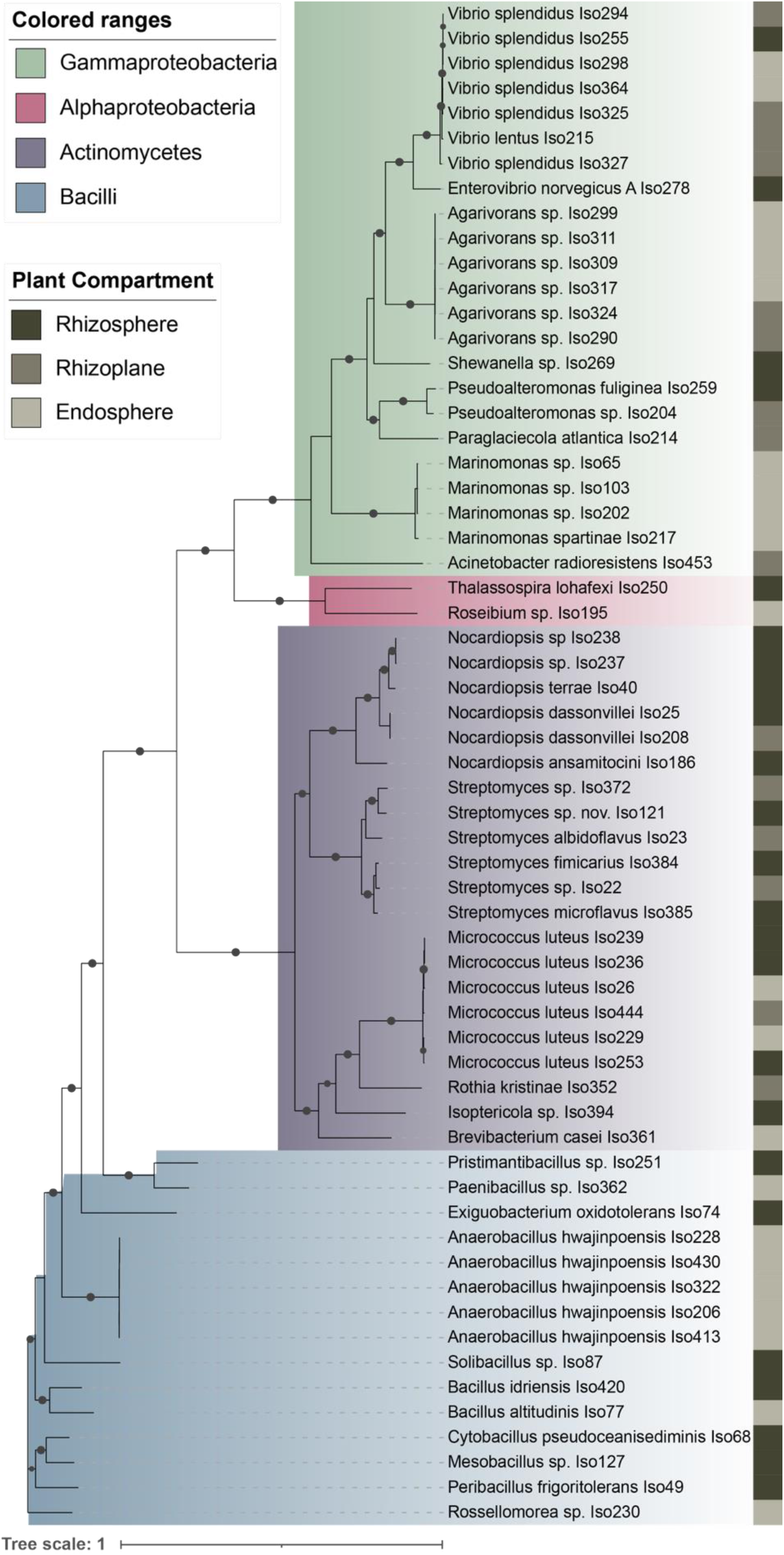
Phylogenomic tree of 61 WGS isolates show the distribution of taxa in the culture collection. The tree was constructed using GToTree with the Bacteria single-copy gene set. Sequences were aligned using MAFFT, and a phylogenetic tree was built with IQ-TREE using the best-fit substitution model and 1000 bootstrap replicates. Taxonomic labels reflect GTDB-tk classification. Color ranges on the tree are colored by phylum while color strip next to the tree indicate which compartment the isolated was collected from. Bootstrap support >95% are shown.

### Genome characteristics of *Z. marina* isolates

All 61 assembled genomes were at least 98% complete and had less then 3% contamination (**Supplementary Table 3**). Genome sizes ranged from 2.38 Mb to 8.41 Mb and had on average of 66.4 ± 28.1% GC content. Of the 61 genomes, 51 were assembled as complete circular chromosomes. Of the ten remaining linear genomes from the remaining six isolates belonged to the genus *Streptomycetes* and contained 1 – 73 contigs, which are all feature common across the genus^63^. Sixteen of the 51 circular genomes also contained accessory plasmids that 2,857 bp to 364,947 bp. The genomes of eight Vibrionaceae isolates (seven *Vibrio* and one *Enterovibrio*) contained two circular chromosomes, consistent with the multi-chromosome architecture common in that family^64^. Two *Pseudoalteromona*s isolates also harbored large accessory replicons (774 kb and 1,036 kb). On average, each genome had a coding density of 86.5 ± 2.2% and contained 2,068–7,592 open reading frames per genome. Finally, we were able to recover between 28-46 unique tRNAs and the 16S rRNA from all 61 genomes (**Supplementary Table 3**).

### Presence of genomic features and in vitro phenotypes in *Z. marina* isolates

Each isolate’s genome was screened for genes involved in nitrogen, sulfur, phosphorus, and phytohormone metabolisms (**Fig. 3, Supplementary Table 4**). Approximately 96% of the isolates contained genes associated in nitrogen mobilization, including those involved in nitrogen fixation (10%), denitrification (51%), DNRA (71%), and C-N bond cleavage (83%). Additionally, 52% of the isolates had the genes for sulfur/thiosulfate oxidation, 88.5% for phosphorus solubilization, and 60.5% for the production of the phytohormone indole-3-acetic acid (IAA) (**Fig. 3c, Supplementary Table 4**). The presence of the genes in the genomes corresponded to the results from our in vitro assays (**Fig. 3b**). Specifically, we found that most of the isolates that tested positive in our assay experiments also had the genes to preform that function (**Fig. 3b,c**, see **Supplementary Text 1** for description of all assay results).

**Fig. 3.**
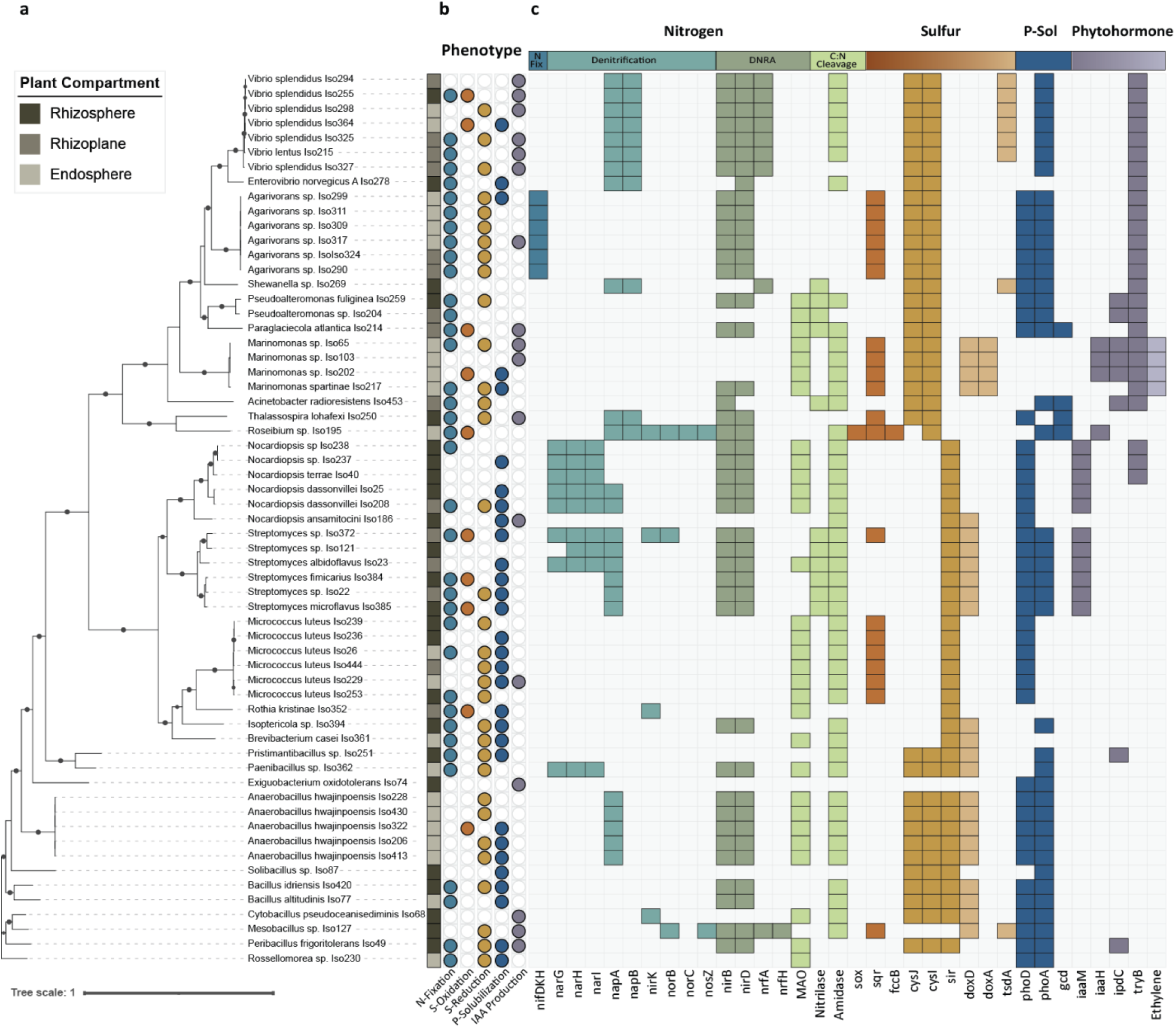
Phenotype assays and genomic content of bacterial isolates from *Z. marina* revealed that each taxa had diverse metabolisms that are predicted to support plant growth. **a,** Phylogenetic tree of the bacterial isolates shows the source compartment (rhizosphere, rhizoplane, or endosphere) of each isolate. Bootstrap support >95% are shown. **b,** Phenotypic assay results for nitrogen fixation, sulfur oxidation, sulfur reduction, IAA production, and phosphate solubilization. Filled circles indicate positive results; open circles indicate negative results. **c,** Presence (filled) or absence (unfilled) of KEGG orthologs associated with PGP pathways, including nitrogen cycling (nitrogen fixation, denitrification, DNRA, C:N bond cleavage), sulfur metabolism (sulfur oxidation, sulfide oxidation, assimilatory sulfate reduction, thiosulfate oxidation), phosphate solubilization, IAA biosynthesis, and ethylene production within each bacterial isolate’s genome.

In our assays, 37 isolates grew on nitrogen-free media, which indicated that they had the ability to mobilize nitrogen to support their growth. However, only six of these isolates had robust growth, while the growth for the remaining 31 isolates was limited. The six isolates the grew well on the media included the *Agarivorans sp.*, which had the complete nitrogenase gene (*nifHDK)* cluster needed for nitrogen fixation. We considered the remaining 31 isolates, where growth was present but minimal, to be nitrogen scavengers. Of these isolates, 29 coded for nitrilase (K01501), amidase *amiE*, and/or monoamine oxidase (*MOA*), which allowed the bacteria to degrade amino acids to ammonia. Two isolates, *Thalassospira lohafexi* and *Vibrio Splendidus*, lacked genes for these enzymes but contained assimilatory nitrite reductase (NirBD), which is responsible for converting nitrite to ammonia.

In our assays, 26 isolates had a visible clearance zone following growth on Pikovskaya media, indicating that they were capable of phosphorus solubilization. These included isolates from the bacterial families Bacillaceae, Micrococcaceae, Streptomycetaceae, Streptosporangiaceae, Brevibacteriaceae, Celerinatantimonadaceae, Cellulomonadaceae, Marinomonadaceae, Paenibacillaceae, Planococcaceae, and Vibrionaceae. Overall, 88.5% of the isolates that tested positive for phosphorus solubilization had either the alkaline phosphatase D (*phoD*), alkaline phosphatase A (*phoA*), or both genes in their genomes. Importantly, the presence of the *phoD* or *phoA* did not guarantee in vitro activity. In our entire collection, 41 isolates carried phosphatase genes (phoD and/or phoA), but only 56% of those tested positive for P-solubilization. This was true even with members of the same genus. For example, only one of the six *Agarivorans sp.* strains tested positive in our phosphorus solubilization assay, despite all the genomes carrying identical sets of alkaline phosphatase genes.

In our culture collection, 41 isolates grew on thiosulfate media, indicating that they either reduced thiosulfate or oxidized sulfur compounds. These isolates were primarily from the Bacillaceae, Celerinatantimonadaceae, and Micrococcaceae, but also included members from several other families. All 41 isolates, except for *Rossellomorea sp.* (Iso230), had either a sulfite reductase [NADPH] flavoprotein alpha and beta-component (*cysJ*, *cysI*) or sulfite reductase ferredoxin (*sir*), all of which convert toxic sulfides into organic sulfur compounds. In our assays, we found that nine isolates acidified the media, indicating that the isolates were oxidizing sulfur in vitro. These isolates include members of the *Micrococcaceae, Streptomycetaceae, Stappiaceae, Bacillaceae, Alteromonadaceae,* and *Vibrionaceae* families. However, only a single *Roseibium sp*. (Iso195) had the complete set of genes needed to oxidize sulfur to sulfates (*soxABCXYZ*). Of the remaining isolates, there were six, including *Anaerobacillus hwajinpoensis* (Iso322), *Streptomyces* sp. (Iso372, Iso384, Iso385) and *Vibrio Splendidus* (Iso255, Iso364) that had the gene for thiosulfate oxidation (*doxD*) present in their genomes. Finally, the remaining two sulfur oxidizing isolates, *Rothia kristinae* (Iso352) and *Paraglaciecola atlantica* (Iso214), lacked conical genes for sulfur oxidation.

Using the Salkowski assay, we observed that 15 isolates tested positive for the production of the phytohormone IAA. Of these isolates, 11 had at least one and as many as three putative IAA biosynthesis genes in their genome. These genes included tryptophan 2-monooxygenase (*iaaM*), indoleacetamide hydrolase (*iaaH*), indolepyruvate decarboxylase (*ipdC*), and aromatic-amino-acid transaminase (*tryB*). These isolates were members of the Micrococcaceae, Bacillaceae, Alteromonadaceae, Celerinatantimonadaceae, Vibrionaceae, and Marinomonadaceae families. Ten isolates from the genera *Paraglaciecola*, *Agarivorans*, and *Vibrio* had the *tryB,* while the two *Marinomonas* isolates contained the genes *iaaH, ipdC,* and *tryB*.

Finally, we screened the isolate genomes for a series of additional genes that are commonly associated with the promotion of plant growth in terrestrial plants (**Supplementary Table T4**). Over 75% of our isolates had one or more genes involved in detoxification of arsenic. For example, 46 isolates had arsenate reductase (*arsC*). Within our isolate collection, 35 of the isolates had geranylgeranyl diphosphate synthase (*GGPS*), which is a terpenoid and gibberellin precursor, a plant signaling molecule and a source of natural products in terrestrial plants^65^. We also identified 25 isolates that had an ACC deaminase (*ACCd*), which metabolizes a precursor to ethylene. Isolates within our collection also encoded type I (n=30), type III (n=2), type IV (n=4) and type VI (n=16) secretion systems. These secretion systems are important because they are often involved in antimicrobial activity, amelioration of biotic stress in plants, and secrete effectors required for the microbial colonization of plant tissues.

### Selection of minimal community of putative PGP bacteria

We applied genome-scale metabolic modeling using metage2metabo^66^ to our 61 cultivable isolates to identify a minimal consortium (MinCom) capable of producing target metabolites relevant to seagrass health. We reasoned that selecting cultivatable members of the seagrass microbiome would allow us to better design a putative PGP synthetic community (MinCom) that we could test in future studies. Using our bottom-up approach, the model identified 32 minimal community solutions, each containing 5 community members. Three members were classified as “essential symbionts”, present in all solutions: *Streptomyces* (Iso23)*, Mesobacillus* (Iso127), and *Roseibium* (Iso195). Each essential member was considered by the model to be the sole predicted producer of one or more of the target metabolites. For example, *Mesobacillus* (Iso127) produced sulfate through sulfide detoxification via tetrathionate hydrolase (EC 3.12.1.B1), *Roseibium* (Iso195) was the only isolate with the genomic capacity to produce nitric oxide, and *Streptomyces* (Iso23) was the sole predicted producer of indole-3-glycerol phosphate (EC 4.1.1.48), a key precursor in the indole/IAA biosynthesis pathway. Two additional isolates, *Peribacillus sp*. (Iso49) and *Streptomyces sp.* (Iso384), were selected as alternative community members because they increased the overall metabolic scope of the MinCom. All five members encoded for genes that support the solubilization of phosphorus and the production of IAA. Interestingly, while individual isolates were able to produce IAA in lab assays, our modeling approach predicted that many of the members exhibited cooperative IAA biosynthesis behavior. For example, the genes involved in the indole-3-pyruvic acid pathway were distributed across the 5-member community. Together, the 5-member MinCom (MinCom-5) was predicted to produce all 30 target metabolites from the defined seed environment (**Supplemental Table 1**).

While the predicted 5-member minimal community was predicted to produce all target metabolites, it did not identify taxa able to fix or scavenge nitrogen. Yet, we know that seagrass growth in many systems is reliant on nitrogen acquisition by the plant^67,68^. In our culture collection, nitrogen provisioning occurred primarily through the catabolism of organic nitrogen sources. However, nitrogen is often growth limiting in many seagrass systems^69^. Therefore, we augmented the initial MinCom-5 with *Agarivorans* sp. (Iso311), as it grew robustly on nitrogen-free media and encodes the complete nitrogenase gene cluster, suggesting it can fix N_2_. Using a community scope analysis within metego2metobo, the final MinCom-6 was predicted to produce 996 metabolites in comparison to 948 metabolites from MinCom-5. These metabolites included the production of the vitamins B1 (thiamine and thiamine pyrophosphate), B2 (riboflavin), B3 (niacin and niacinamide), B5 (4-phosphopantothenate), B6 (pyridoxal, pyridoxamine, and pyridoxine), B9 (dihydrofolate), and B12 (adenosylcobalamin), which are important co-factors in plant growth^70^. Production of these vitamins was largely redundant across community members, except for niacinamide, which was independently produced by *Peribacillus* (Iso49).

## DISCUSSION

In our study, we were able to isolate 201 putative PGP bacteria from *Z. marina’s* rhizosphere, rhizoplane and endosphere. In doing so, we significantly expanded the number of bacterial species isolated from *Z. marina* and expanded current bacterial genome databases^71,72^. Many of these isolates recovered in our study tested positive for putative beneficial traits in our physiological assays and genomic analyses. Specifically, putative PGP bacteria were able to mobilize nitrogen, assimilate sulfur, solubilize phosphate and produce phytohormones. As in terrestrial system^73–76^, these bacterial traits support plant growth and metabolism by providing seagrasses with key nutrients required to build biomass^16^. Finally, our metabolic modeling approach identified a minimal bacterial community whose members exhibited complementary metabolism reflected these traits. Our proposed MinCom-6 informs current conservation efforts aimed at leveraging wildlife probiotic approaches to conserve and restore seagrass ecosystems^8^.

### Plant-derived media supported cultivation of diverse bacterial taxa

Using our PB cultivation media, we were able to successfully isolate members of multiple groups that are rarely cultivated^71,77^. For example, we expanded the current number of bacteria isolates cultivated from seagrasses to include member of Bacillaceae, Nocardiopsaceae, Rossellomoreaceae and Stappiaceae families. Furthermore, there were nine species in our current culture collection, represented by 17 isolates, that were not represented in the GTDB. This suggests that these nine species, which include members of *Bacillus, Isoptericola, Shewanella, Solibacillus, Roseibium, Streptomyces, Nocardiopsis, Rossellomorea,* and *Agarivorans* genera are novel taxa. In terrestrial systems, members of the *Bacillus* and *Streptomyces* repeatedly promote the growth of a broad diversity of plants^78^. Furthermore, bacteria isolated from saline environments facilitate plant response to drought conditions^79^. For example, *Isoptericola* isolated from mangrove sediments promoted growth of *Arabidopsis* under saline conditions^80^. Finally, we were able to isolate two species from the Rhodobacteracea family, which was the most relatively abundant family within the 16S rRNA amplicon libraries across seagrass compartments. The Rhodobacteracea, which is one of the nine most abundant lineages in marine ecosystems, includes taxa that occur in marine sediments and within the phycosphere of marine algae, and are involved in a number of metabolic processes^81,82^. Many of the genera in this group are challenging to cultivate^81^, however our culture collection included species from the *Pseudosulfitobacter* and *Roseicitreum* within the Rhodobacteracea family. Together, these results highlight the importance of leveraging host-derived media, which likely contains key host factors^24^, to support cultivation of difficult to isolate bacteria.

Not surprisingly, our aerobic cultivation approach did not isolate bacterial members from strict anaerobic groups. Yet, cultivating anaerobic bacteria from seagrasses is likely important given that sulfate-reducing families Desulfocapsaceae and Desulfobacteraceae comprised 9.69% of relative abundance in our 16S rRNA amplicon dataset, which are obligate anaerobic taxa^24,83,84^. Moreover, these taxa are likely ecologically relevant to seagrasses and sediment biogeochemical cycles as they play an important role in reducing sulfate to form sulfide in marine systems^85,86^. In rice, which also occupy anoxic soils, the reduction of sulfate into H_2_S by members of the Desulfocapsaceae increases the content of abscisic acid and betaines in saline-alkali soils which is tied to increased plant production^87^. Consequently, it is possible that these taxa, while producing toxic sulfides, might also be linked to increased plant productivity. Future cultivation efforts should consider combining our PB media approach with anaerobic cultivation conditions to isolate members of these missing families to support our understanding of these taxa in the promotion of seagrass growth.

### Putative PGP bacteria support *Z. marina’s* acquisition of nitrogen and phosphorous

In many coastal ecosystems, including in *Z. marina* meadows, the availability of nitrogen and phosphorous limits seagrass growth and reproduction^88–90^. The availability of these two important macronutrients is often dependent on abiotic and biotic factors that are driven by sediment conditions, light, seasonality, temperature, pH and oxygen availability^88,91,92^. Experimental manipulation of nitrogen and phosphorous levels directly increases seagrass biomass^89^, suggesting both nutrients are critical towards enhancing seagrass growth. Consequently, by identifying and isolating bacteria from *Z. marina* that can support nitrogen and phosphorous acquisition, it is possible to experimentally test if these bacterial taxa can support seagrass acquisition of these limiting nutrients.

In our study, we identified 37 taxa in our sequenced-culture collection that were able to either liberate ammonia from nitrogenous compounds or fix atmosphere N_2_. *Z. marina* and other seagrasses primarily take up nitrogen in the form of ammonium through their root system^93^. Many of the bacteria isolated from *Z. marina’s* root endosphere, rhizoplane and rhizosphere had either a nitrilase, amidase and monoamine oxidase enzyme, which would allow them to catabolize amino acids or proteins to produce ammonia that is then ionized into ammonium within the plant’s sediment pore waters. This trait is likely common across seagrass meadows as some members of the microbiome from *Posidonia oceanica* also catabolize amino acids to liberate ammonium that is then available for plant use^67^.

Interestingly, both *Z. marina* and *P. oceanica* also host closely related N-fixing bacteria within their root tissues. The six strains of *Agarivorans sp.* isolated from *Z. marina* had robust growth on N-free media and encoded the complete nitrogenase gene cassette (*nifHDK*) needed for nitrogen fixation, suggesting these are likely N_2_ fixing bacteria. The *Agarivorans* species belong to the same family as *Candidatus* Celerinatantimonas neptuna, which is the N_2_-fixing symbiont within the root of *P. oceanica* ^21,94^. Given that both *Z. marina* and *P. oceanica* host diverse taxa that support N-assimilation through multiple pathways within their root tissues, the microbial enhanced acquisition of nitrogen may be a key trait in supporting robust seagrass growth.

Nitrogen metabolism within *Z. marina’s* root endosphere, rhizoplane and rhizopshere may be contingent on metabolic flexibility of individual community members and microbial cross-feeding. This is likely due to the sharp transition zones in oxic and anoxic regions within the root to sediment interface as oxygen leaked from the roots is rapidly metabolized by the microbial communities living within the rhizosphere^19,91^,. The resulting redox gradients select for metabolic flexibility needed to transform nitrogenous compounds into ammonium. In our isolation collection, over 80% of all isolates that encoded for the conversion of dissimilatory nitrite to ammonia (*nirBD*), a reaction which occurs under anoxic conditions, also had an amidase (*amiE*), which converts amides to ammonia regardless of the oxygen concentrations. Furthermore, no one isolate contained the complete denitrification and DNRA pathway. Rather, the capacity for both processes occurred across multiple members of the microbiome, a finding that is consistent with recent results showing that denitrification and DNRA are community traits^95^. Our findings suggest that PGP consortia design for seagrass restoration and conservation should consider identifying community members with complementary nitrogen cycling capabilities rather than relying on single isolates with complete pathways.

Seagrasses, including *Z. marina*, take up inorganic phosphorus in the form of phosphate through their roots using phosphate transporters^69^. Because seagrass sediments typically contain high concentrations of organic matter, inorganic phosphorus is typically locked away within the organic matter itself^92,96^. To liberate phosphate, seagrasses secrete organic acids to stimulate the activity of microorganisms that have the capacity to mineralize phosphorus^87^. Of the bacteria that underwent whole genome sequencing, 26 isolates were able to mineralize phosphorus and liberate phosphate using alkaline phosphatases (*phoD, phoA*), which supports the ability of *Z. marina* to acquire phosphorus in situ. Not all isolates that had the genomic ability to produce phosphate tested positive in the phosphorus solubilization assays. For example, some strains had identical sets of phosphatase genes yielded different P-solubilization outcomes, likely owing to the fact that our lab assays do not replicate the complex geochemical environment of the rhizosphere^97,98^. Seagrasses exhibit temporal secretion of oxygen which creates spatially variable pH gradients within the rhizosphere, likely impacting pH driven release of sediment bound phosophorus^99^. Consequently, it is possible that differences in our assay results reflect physiological adaptions of different bacterial taxa living in varying niches of *Z. marina* root and rhizosphere system. To overcome variation in the capacity of *Z. marina’s* microbiome to liberate phosphorus to support plant growth, it is critical to identify which microbial taxa can transform phosphorus into phosphate.

### Putative PGP bacteria detoxify sulfides in anoxic sediments

In our culture collection, 41 isolates grew on thiosulfate media, the majority of which were able to reduce sulfite using genes (*cysJ, cysI, sir*) that convert toxic sulfides into organic sulfur compounds. Our result suggest that sulfur detoxification is a widespread trait across members in our collection. In the context of identifying putative PGP bacteria for seagrasses, our findings are important because *Z. marina* root tissues are susceptible to sulfide intrusions as they occur in highly reduced sediments^100^. Sulfide intrusions are toxic to seagrasses and their accumulation can lead to meadow dieback^101^. To protect their root tissues, *Z. marina* and other seagrasses release oxygen from their root tips to generate protective oxic zones that work to recruit sulfur-oxidizing bacteria^19,91,102^.

In our culture collection, we identified nine isolates that were able to oxidize sulfur. However, only *Roseibium sp.* (Iso195) has the complete Sox pathway for sulfur oxidation pathway (*soxXYZABCD*). *Roseibium* and other members of the Rhodobacteraceae are often implemented in sulfur oxidation and play a key role in DMSP cycling^82^ and oxidate stress responses in marine invertebrates and red algae^103,104^. Here, we show that *Roseisbium* sp. also associate with seagrass roots, likely plays a critical role in sulfur oxidation. In addition to *Roseibium*, our isolate collection included six taxa that were able to oxidize sulfur using thiosulfate dehydrogenase (*doxA, doxD*) and tetrathionate hydrolase (*TetH*) pathways. Tetrathionate hydrolase has been characterized primarily in acidophilic sulfur-oxidizing bacteria, making its presence in marine, plant-associated bacteria poorly understood^105^. Only one isolate in our collection, *Mesobacillus sp*. (Iso127) encoded the *TetH* gene. In this pathway, tetrathionate is hydrolyzed to thiosulfate, elemental sulfur, and sulfate^105^. Tetrathionate is present in marine sediments but rarely detected because concentrations are below detection limits, suggesting active microbial cycling^106^. The presence of *TetH* in *Mesobacillus sp*. suggests this pathway may contribute to sulfur cycling in *Z. marina* sediments, potentially playing a role in processing sulfur intermediates. By isolating multiple sulfur cycling bacteria from *Z. marina*, future studies can leverage genetic manipulation experiments to determine the mechanism that these taxa use to reduced sulfide intrusions in the plant’s tissues.

### Putative PGP bacteria produce phytohormones

In terrestrial systems, the presence of the phytohormone IAA stimulates root growth and cell elongation, while providing defense against pathogens^75^. It is also a necessary trait for the bacterial colonization of plant root systems^22,107^. In our culture collection, 18% of the isolates were able to produce IAA. This might allow seagrasses and their microbiome to engage in a mutually beneficial feedback loop: IAA produced by bacteria induces plant immune responses that enhance bacterial colonization and pathogen inhibition^107^. The genome of *Z. marina* encodes the auxin response factor that would bind IAA^108^ and prior metatranscriptomics data indicates that microbial taxa living within the *Z. marina* rhizosphere actively convert tryptophan to IAA in the field^18^. Our work expands on these findings by showing that a broad diversity of bacteria species and metabolic pathways lead to the biosynthesis of IAA.

On land, plants release tryptophan through their roots to their rhizosphere sediments, providing the initial substrate for the production of IAA^109^. Similar patterns likely occur in seagrass meadows, as tryptophan is a common root exudate in seagrasses^110,111^. This could modulate microbial activity by recruiting bacteria that consume tryptophan and convert it to IAA or similar compounds. In our study, isolates encoded enzymes involved either in the indole-3-acetamide (IAM) pathway and the indole-3-pyruvic acid pathway (IPA). The IAM pathway uses tryptophan monooxygenase (*iaaM*) to convert tryptophan to IAM followed by hydrolysis IAM to IAA by indole-3-acetamide hydrolase (*iaaH*)^112^. In the IPA pathway, tryptophan is first converted to IPA by aminotransferase (*tryB*), then pyruvate decarboxylase (*ipdC*) converts IPA to indole-3-acetaldehyde, which is oxidized to IAA by aldehyde dehydrogenases^112^.

We found that IAA synthesis pathways were prevalent in the isolate collection but often incomplete within individual taxa. For example, *Streptomyces* sp. and *Nocardiopsis* sp. possessed *iaaM* but lacked *iaaH,* and therefore cannot complete the final hydrolysis step to transform IAM to IAA. Consistent with our genomic prediction, these strains tested negative in our IAA production assays. Taxa with partial IAA pathways may produce and release IAA intermediates (e.g., indole-3-acetamide) or precursors that community members convert to IAA. Supporting this idea, *Roseibium sp.* (Iso195) in our culture collection possessed an *iaaH*, which would enable it to transform IAM produced by *Streptomyces* or *Nocardiopsis* into IAA, facilitating community-level phytohormone production. This mechanism of community cooperation may explain why *Streptomyces* species are commonly included in plant growth-promoting consortia^22^.

### MinCom-6 is predicted to support plant growth through nutrient provisioning to *Z. marina*

Genome-scale metabolic modeling integrates expected metabolic reactions of individual host organisms and their microbial communities, enabling prediction of the minimum set of microbial taxa needed to produce target metabolites given specific nutritional constraints^113^. We applied this approach to predict which members of our isolate collection produced key metabolites predicted to promote plant growth, including amino acids, vitamins, and phytohormones. Furthermore, we were able to identify functional redundancies across members of the microbiome while identifying likely cross-feeding interactions across community members^62,114^. This approach has helped to identify synthetic communities that promoted plant growth in sorghum in controlled and native field settings^10^ and in wheat when grown in contaminated soilds^115^. Consequently, we reasoned that we could leverage this approach to identify a synthetic community of microbial taxa that could be tested in future studies to support the growth of seagrasses.

By combining our modeling approach and in vitro assays, we identified six members of our culture collection (MinCom-6) that are predicted, as a community, to support *Z. marina* growth. Three of these taxa, including *Streptomyces sp*. (Iso23), *Roseibuim sp.* (Iso195), and *Mesobacillus sp.* (Iso127) were considered essential in the model as they were predicted to be the sole producers of one or more metabolites. For example, *Streptomyces sp*. (Iso23) was the only bacteria in our collection that was predicted to produce indole-3-glycerol phosphate, which is an important intermediate in the tryptophan metabolism pathway and serves as a precursor to IAA^112^. *Roseibium sp.* (Iso195) was predicted to be the sole producer of nitric oxide, a root growth signaling molecule^116^ and key metabolite in the DNRA pathway^117^. Nitric oxide also plays a role in alleviating damage of salinity stress in *Z. marina*^118^. Finally, *Mesobacillus sp* (Iso127) was the only bacteria in our collection that produces sulfate via the tetrathionate hydrolase pathway. Two other bacterial taxa, *Peribacillus* (Iso49) and *Streptomyces sp*. (Iso384) were also included as community members in the model due to their ability to expand the metabolic scope of the MinCom. Finally, because nitrogen availability fluctuates with the environment (season, light availability, temperature, and oxygen concentrations)^20,91^, we chose to include the N_2_-fixing *Agarivorans* sp. (Iso311) within MinCom-6. This supports the capacity of *Z. marina* to acquire nitrogen through diverse metabolic pathways.

## CONCLUSIONS

In our study, we expand the current culture collections and genomic diversity of bacterial taxa that are predicted to promote seagrass growth. In doing so, we established pure cultures of bacterial taxa, including the addition of novel species to current genomic databases, which can provide insights into bacterial functional potential. By integrating our novel cultivation approach with genomics, phenotypic assays and metabolic modeling, we were able to select a suite of bacterial isolates that serve as a launching point in the development of PGP bacteria for seagrass meadows. Our framework bridges the divide between sequence-base discovery and actionable microbial solutions for supporting seagrass restoration and conservation efforts

We focused on bacteria that exhibited traits known to promote seagrass growth, including nitrogen mobilization, phosphate solubilization, sulfide detoxification and the production of phytohormones. This generated an extensive list of putative PGP bacteria for seagrasses. Given that globally seagrasses are in decline, our study is timely as it provides a new research avenue for identifying PGP bacteria that work to support meadow expansion. Future research requires careful experiments to test the efficacy of delivering MinCom-6 or similar consortia to seagrasses root and rhizosphere systems. Considering that PGP bacteria promote the productivity of terrestrial plants^119^, we predict that the taxa identified here will support the application and implementation of wildlife probiotics in seagrass meadows.

## Supporting information

Graphical abstract

## SUPPLEMENATRY INFORMATION

### AVAILABILITY OF DATA AND MATERIALS

Raw data used in this study are available in the NCBI Sequence Read Archive (SRA), accession numbers PRJNA1260012. The accession number for each assembled genome is provided in **Supplementary Table 3**. https://doi.org/10.5281/zenodo.19102533 and https://github.com/DBrache/ZM_PGPR

## ACKNOWLEDGEMENTS

We thank Drs. Anya Brown and Jay Stachowicz, and their labs at UC Davis Bodega Marine Laboratory for use of lab, field, and storage spaces, and Serina Moheed for logistical support. We thank Dr. Jonathan Eisen and the Eisen lab for guidance on Nanopore 16S rRNA amplicon bioinformatics, and Drs. Stachowicz, Eisen and Gina Chaput for advice throughout this project.

## Abbreviations

IAA: Indole-3-Acetic Acid
DNRA: dissimilatory nitrate reduction to ammonium
PGP: Plant growth promoting bacteria
PB: plant-based media
SASW: sterile artificial seawater
MB: Marine Broth
ONT: Oxford nanopore technology
ASV: Amplicon sequence variant
M2M: metage2metabo pipeline

## AUTHOR INFORMATION

### Corresponding authors

Correspondence to Diane Brache-Smith and Maggie Sogin

### Contributions

DB and MS designed the study. DB and JB conducted seagrass sample collection and initial bacterial culturing. DB, JB, and SM isolated and assayed the culture collection. DB conducted DNA extractions, library preparations, bacterial genome sequencing, and performed bioinformatics, analysis, and visualization. DB wrote the first draft; MS reviewed and edited subsequent drafts. All the authors read and approved the final manuscript.

