## Supplementary figures and images for "Identification of bacterial candidates that promote the growth of the seagrass *Zostera marina*"

### Graphical abstract

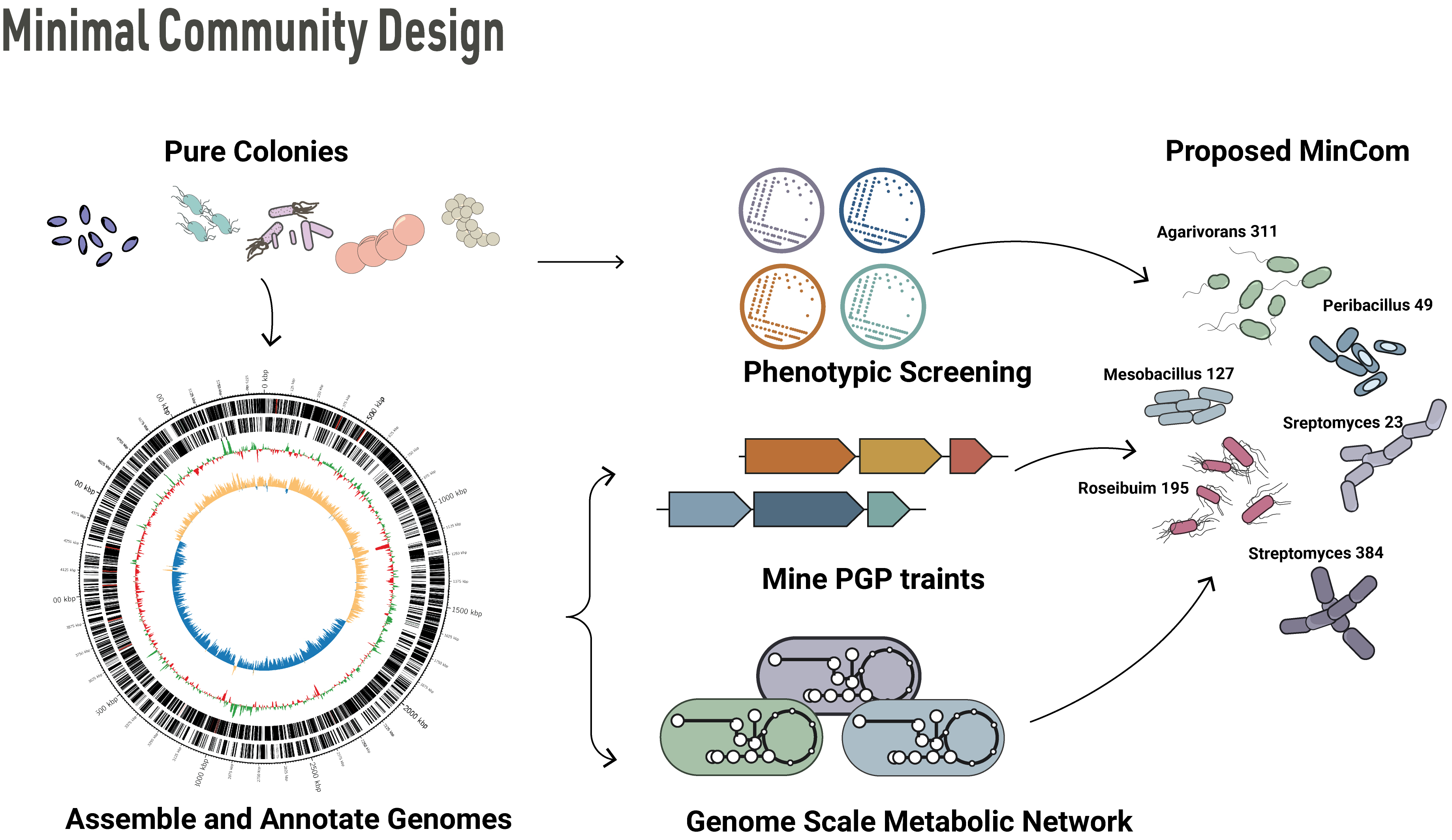

### Supplemental Figure 1

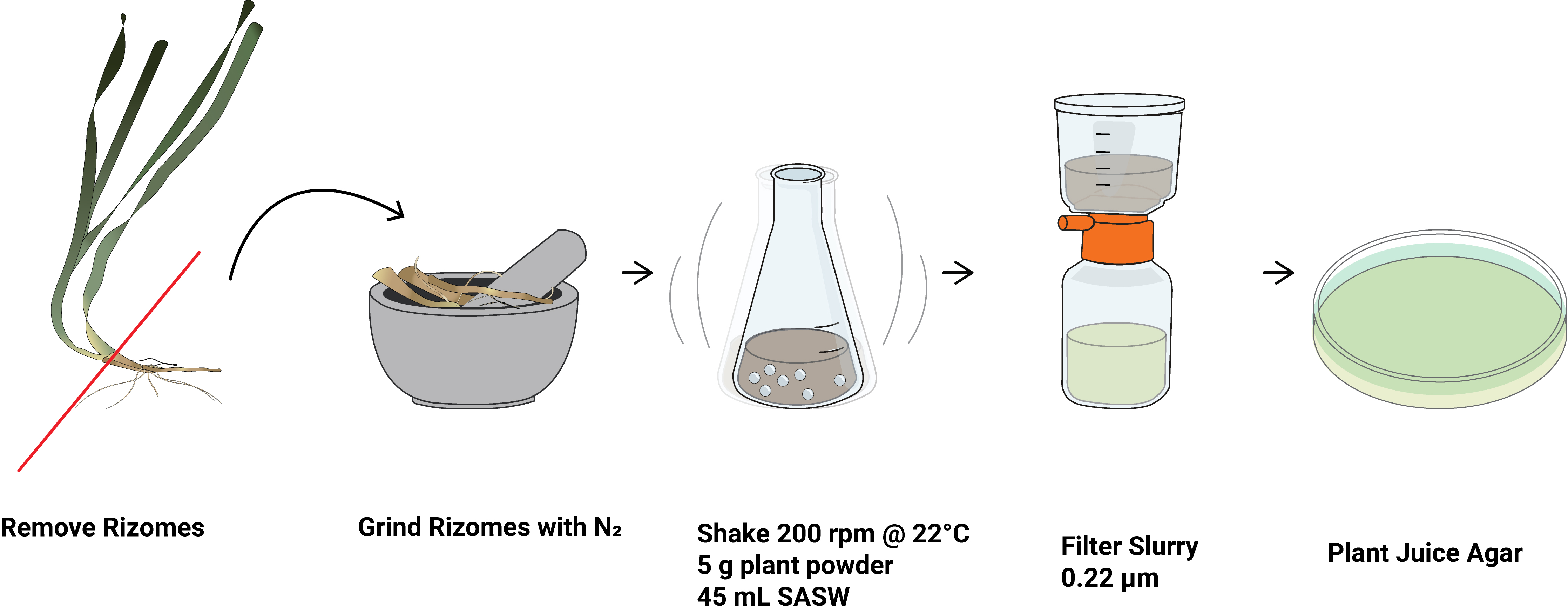

### Supplemental Figure 2

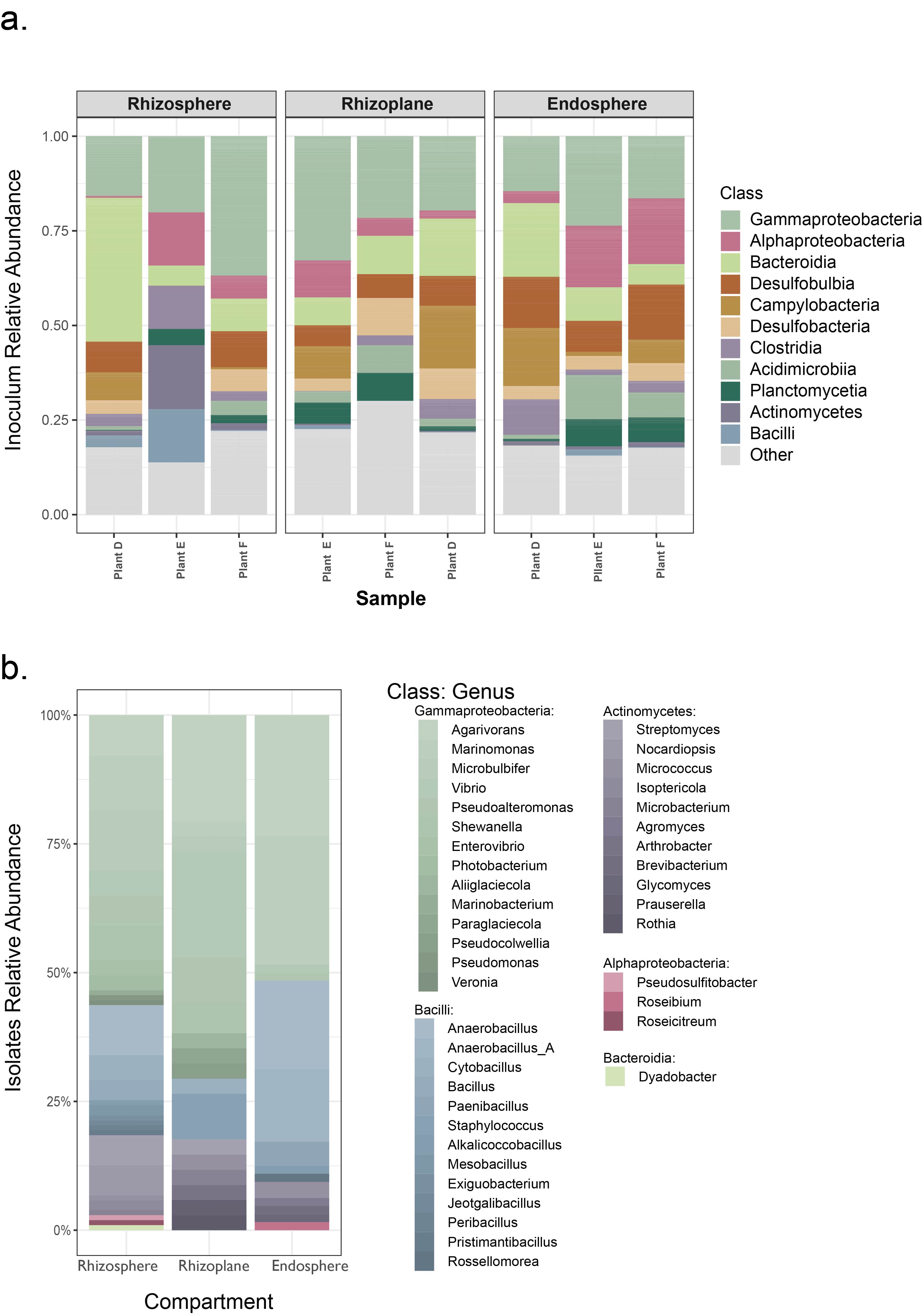

### Supplemental Figure 3

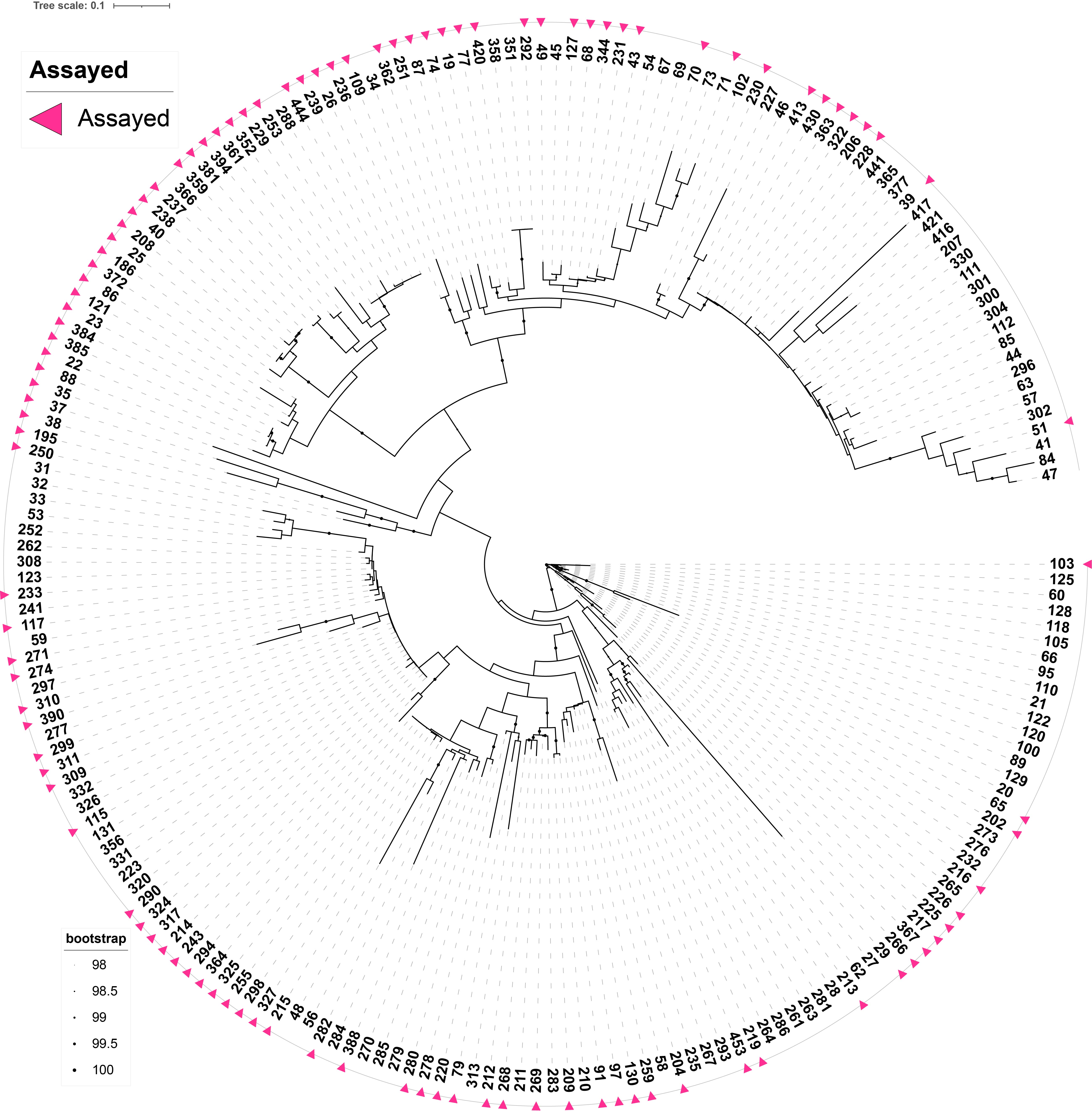

### Supplemental Figure 4

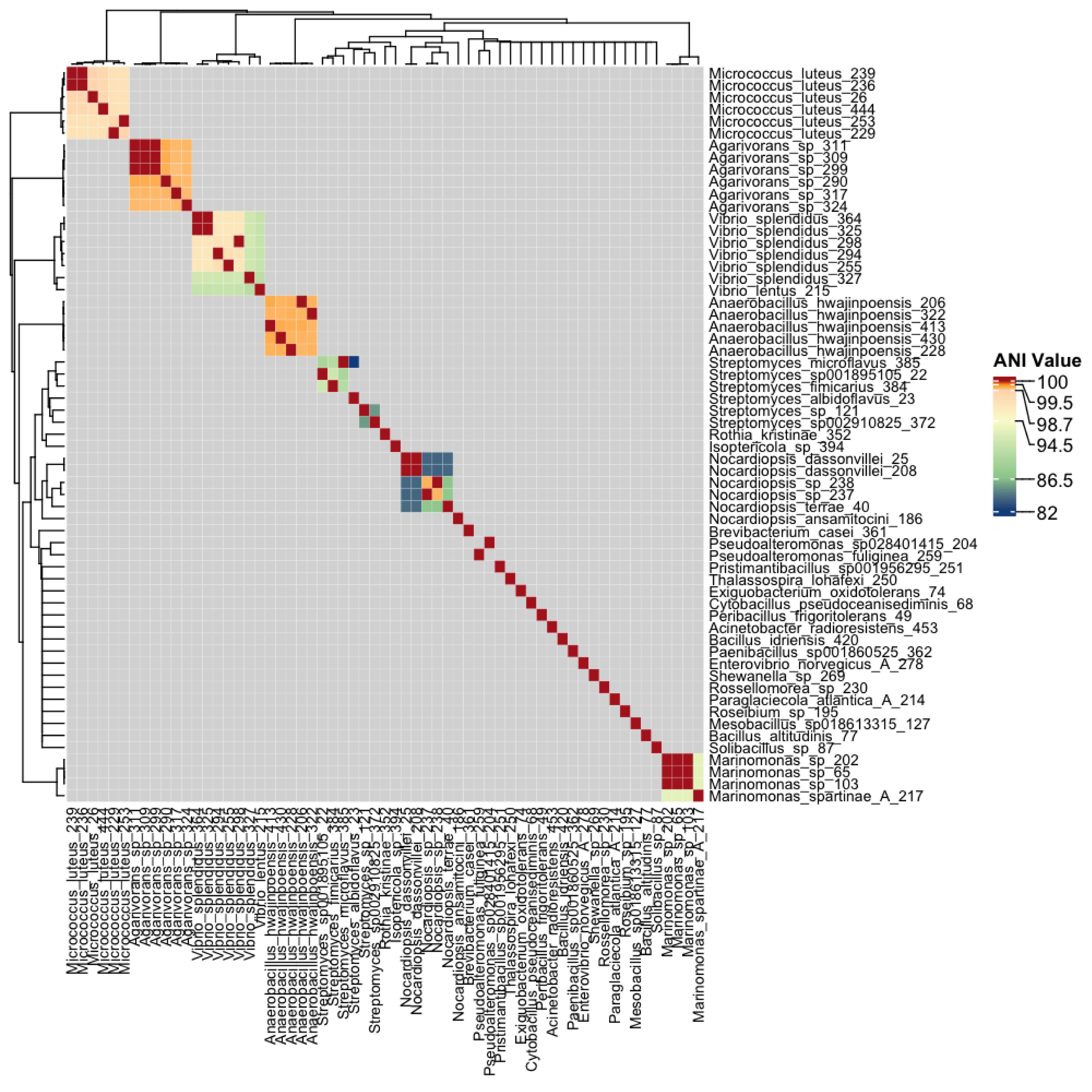
